# Development of whole-porcine monoclonal antibodies with potent neutralization activity against classical swine fever virus (CSFV) from single B cells

**DOI:** 10.1101/389361

**Authors:** Haisi Dong, Dongmei Lv, Ang Su, Lerong Ma, Jianwei Dong, Nannan Guo, Linzhu Ren, Huping Jiao, Daxin Pang, Hongsheng Ouyang

## Abstract

Classical swine fever (CSF) is a highly contagious swine disease found worldwide that has caused devastating economic losses. However, there are few efficacious mAbs against the CSF virus (CSFV) that can be used for treatment because most mAbs against CSFV are derived from mouse hybridoma cells and these murine mAbs have disadvantages of inefficient effector functions elicitations and high immunogenicity *in vivo*. Accordingly, we characterized whole-porcine anti-CSFV neutralizing mAbs (NAbs) isolated directly from single B cells sorted from a CSFV-vaccinated pig using the fluoresceinated conserved linear neutralizing epitope of the CSFV E2 protein and fluorophore conjugated goat anti-pig IgG. Immunoglobulin (Ig) genes were isolated via nested PCR, and two porcine mAbs termed HK24 and HK44 were produced. We determined that these mAbs can bind to E2 protein and recognize sites within this major antigenic epitope. In addition, we found that mAbs HK24 and HK44 exhibit potent neutralizing activity against CSFV, and they can protect PK-15 cells from infections *in vitro* with potent IC_50_ values of 9.3 μg/ml and 0.62 μg/ml, respectively. Notably, we demonstrated that these two mAbs can be used as novel reagents for detecting virus infection. These data suggest that our results not only provide a method for efficiently obtaining mAbs against CSFV but also offer promising mAb candidates for development of antibody-based diagnostic and antiviral agents.

**Importance:** Neutralizing monoclonal antibodies (NAbs) can prevent and may slow the spread of virus infection. The discovery of NAbs that recognize classical swine fever virus (CSFV) necessitates new technologies because the NAbs produced by immunization and hybridoma technology could not be transferred to *in vivo* research. Multiple full-length human therapeutic antibodies have been produced via single-cell polymerase chain reactions but whole-porcine NAbs for CSFV have not been generated. In this study, two whole-porcine mAbs, named HK24 and HK44, were isolated from epitope-specific single B cells. We demonstrate that these two mAbs have potent neutralizing activity against CSFV and can protect cells against viral infection. Therefore, they may facilitate the development of vaccines or antiviral drugs that offer the advantages of stability and low immunogenicity.

## Introduction

Classical swine fever (CSF) caused by classical swine fever virus (CSFV) is a highly contagious and fatal viral swine disease that remains a serious problem for the pork industry (1–3). CSFV is an enveloped virus with positive-sense RNA and belongs to the genus *Pestivirus* of the family *Flaviviridae* (4). The mature CSFV virion contains twelve viral proteins, N^pro^, C, E^rns^, E1, E2, p7, NS2, NS3, NS4A, NS4B, NS5A, and NS5B (5). E2 is a protein with multiple functions that can form a dimer with E1 and mediate virion entry into target cells; additionally, it is the major antigen for the production of neutralizing antibodies that protect the body from the virus. The ability of an anti-E2 antibody to neutralize CSFV has previously been characterized in detail (6, 7). The E2 protein consists of four antigen domains, A, B, C, and D, at the N-terminus with a total of 177 amino acids, ranging from amino acids 690 to 866. These domains contain some conformational epitopes and linear epitopes that play critical roles in inducing neutralizing antibodies. Comparison of the different epitopes of the E2 protein has demonstrated that the linear epitope CTAVSPTTLRTEVVK, found between amino acids 828 and 842, a region recognized by the monoclonal antibody WH303, is strongly conserved in different CSFV strains but is highly divergent among bovine virus diarrhoea virus (BVDV) and border disease virus (BDV) strains (8, 9).

NAbs that are capable of directly neutralizing most strains of a given highly antigenic variable pathogen have attracted considerable attention because they can used to treat some of the difficult infectious diseases encountered in modern medicine and have potential for development of passive immunotherapy treatment or vaccine reagents (10). In addition, given their potential antiviral effects, NAbs also play an essential role in studying the structure-function properties of infectious agents and related pathogenesis. However, mAbs of mouse origin that are produced by hybridoma technology cannot be applied in *in vivo* research because they can induce the production of anti-mouse antibodies *in vivo*, leading to a short half-life (11–13). Other traditional methods of generating mAbs, such as Epstein-Barr virus-transformation and phage-display libraries, are limited by being unstable, time consuming or inefficient (14, 15). With the advent of single-cell reverse transcription PCR (RT-PCR) technologies, the amplification of full-length immunoglobulin gene (Ig) fragments from single B cells by utilizing nested PCR has allowed the bypass of many of these limitations and this technology has been shown to provide a versatile tools for generation of new mAbs; this is an important advancement in mAbs production (16, 17). Prior to the amplification of Ig-encoding genes, antigen-specific memory B cells must be stained with fluorescently-labelled antigen baits and then sorted. Several human mAbs against HIV, ZIKV and Ebola virus have been obtained through this technology (18–20). However, the production of whole-porcine mAbs against the E2 protein of CSFV using linear specific-epitopes has not been reported. Whole-porcine neutralizing antibodies usually exhibit the lowest immunogenicity, longer half-lives and have a great potential for superior applications in *in vivo* (21, 22).

In this study, we report the acquisition of the whole-porcine mAbs HK24 and HK44. These mAbs were isolated from single B cells of a vaccinated pig using the fluoresceinated epitope CTAVSPTTLRTEVVK and fluorescein isothiocyanate (FITC)-labelled goat anti-pig IgG via fluorescent-activated cell sorting (FACS) (23). Their ability to significantly neutralize CSFV was identified through a panel of assays. We also confirmed that these mAbs demonstrate significant value for the serological diagnosis of CSFV infection. Together, the porcine mAbs HK24 and HK44 show potential as candidates for immunotherapy or diagnostic reagents.

## Results

### Isolation of single B cells from a vaccinated pig and mAb generation

Individual pigs were immunized with an attenuated vaccine strain of CSFV to generate a spectrum of antibody responses ranging from low to high levels of blocking, which was determined using the Classical Swine Fever Virus Antibody Test Kit (IDEXX, Switzerland); unvaccinated pigs served as negative controls (**Fig. 1A**). As shown in **Fig. 1B**, pigs #3748 and #3757 exhibited a high blocking rate; therefore, blood samples from pig #3748 were used to isolate single B cells that were stained with FITC-conjugated anti-pig IgG and 5-TAMRA-conjugated epitope-76 via FACS. Epitope-76 is strongly conserved in different CSFV strains but is highly divergent among the BVDV and BDV strains, as shown in **Fig. 1C**. As shown in **Fig. D**, epitope-specific IgG+ B cells constituted approximately 0.39% of the cell population. Full-length heavy- and light-chain antibody genes were amplified from the cDNA of single purified B cells via nested PCR and were sequenced. All nine antibodies were obtained from one hundred single epitope-specific IgG^+^ B cells. Nucleotide sequence analysis of these genes demonstrated that they could be divided into four different groups. Some of these antibodies shared the same *IgH* and *IgL* genes, and most clones were somatic variants of the IGHV1-4*02 and IGKV1-11*01/IGKV2-10*02 germline genes (**Fig. 1E**). This result was similar to that obtained in a study of ZIKV (20).

**Figure 1.**
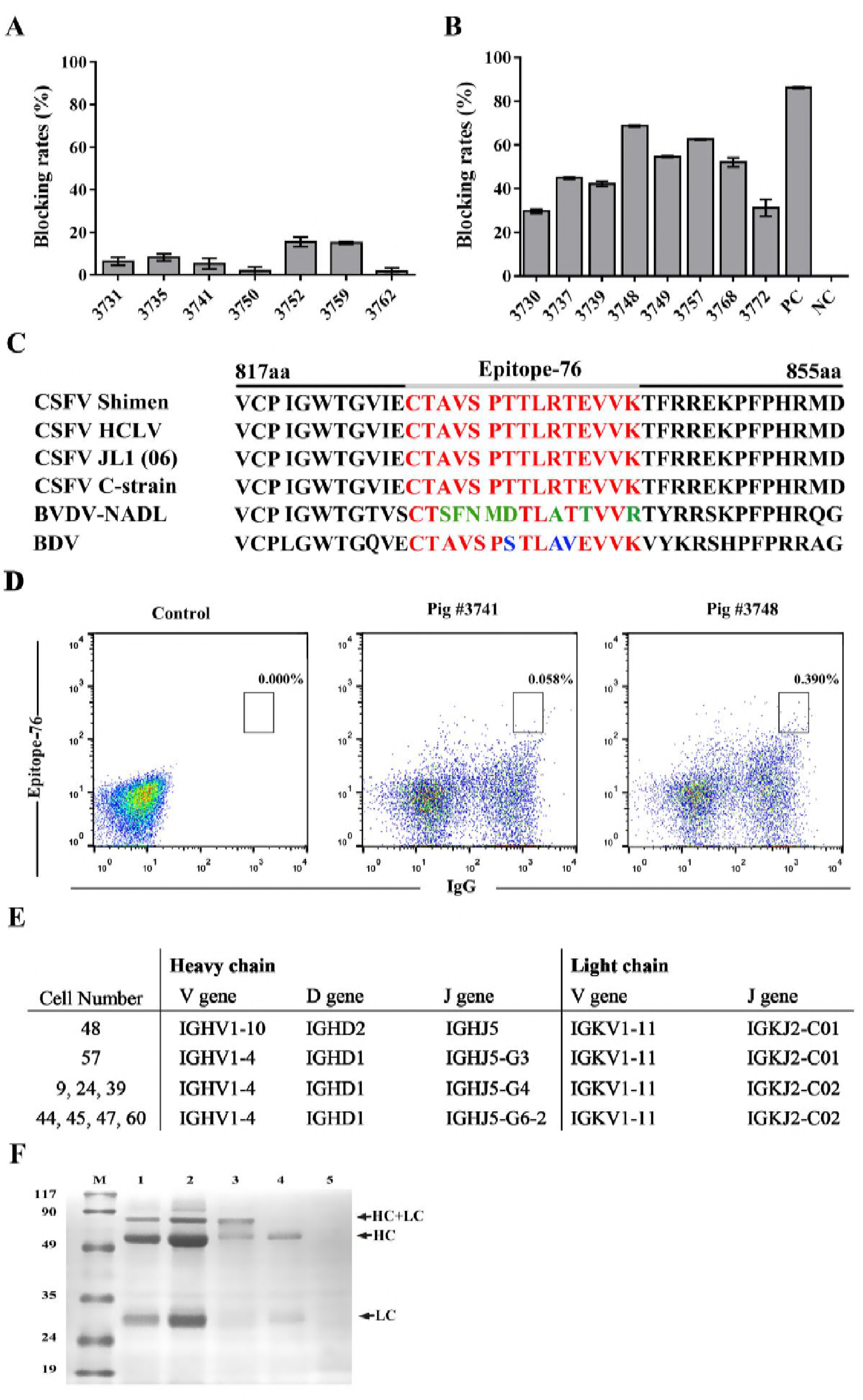
Isolation of epitope-specific antibodies. (A and B) Detection of CSFV antibodies was performed using a CSFV antibody test kit (positive blocking rate ≥ 40%, negative blocking rate ≤ 30%). (A) Blocking rate of serum samples isolated from unvaccinated pigs. (B) Blocking rate of serum samples isolated from individuals immunized with attenuated vaccine strain of CSFV. (C) Sequence alignment between E2 from four different CSFV strains, one BVDV strain and one BDV strain. The conserved amino acid residues are highlighted in red, and the somatically mutated residues are highlighted in green (BVDV stains) and blue (BDV strains). (D) Frequency of epitope-specific, IgG+ memory B cells in the peripheral blood of pig #3748. An unvaccinated pig, #3741, was used as the negative control. Flow cytometry plots display the percentage of IgG^+^ B cells that bound to FITC-conjugated epitope-76 bait. (E) List of V (D) J segments of antibody genes that we isolated and germline (GL) assignment which were derived using the international ImMunoGeneTics information system (IMGT). (F) Analysis of the expression of the mAbs HK24 and HK44 via SDS-PAGE. Samples were resolved by reducing 12% SDS-PAGE, followed by staining with Coomassie blue. Lane M, marker protein; lane 1, porcine IgGs 8 μg; lane 2, porcine IgGs 20 μg; lane 3, mAb HK24; lane 4, mAb HK44; lane 5, supernatant of control HEK293T cells. The black arrows on the right indicate the antibody heavy chains (HC) and light chains (LC).

In summary, we deduced that individuals with high serologic neutralizing titres against the same epitopes of CSFV may express isotype antibodies. We also found that the mAbs HK24 and HK44 each had a long complementarity determining 3 region of anibody heavy-chain (CDR H3) loop composed of 17 and 21 amino acids, respectively. The antibody gene pairs HK24 *IgH, IgK* and HK44 *IgH, IgK* were selected to produce mAbs, which were identified via SDS-PAGE. Polyclonal antibody IgGs were isolated from porcine serum by protein A were served as a positive control (**Fig. 1F**).

### The mAbs HK24 and HK44 can bind to specific epitopes of CSFV

To map the epitopes of the mAbs HK24 and HK44, we first checked whether these mAbs react with epitope-76. For this purpose, HEK293T cells were used for the transient expression of mAb HK24 or HK44, and the binding activity of the mAbs HK24 and HK44 to FITC-conjugated epitope-76 (20 μg/ml) was detected and analysed through microscopy. As shown in **Fig. 2A**, there was a high level of green fluorescence in cells transfected with the expression vector for either HK24 or HK44, and cells without expression vectors showed no green fluorescence. This result indicated that HK24 and HK44 could bind to the conserved epitope-76 with a higher affinity. This distinctive feature of the mAbs HK24 and HK44 was further confirmed by flow cytometric analysis, as shown in **Fig. 2B**. FACS results indicated that a higher percentage of HEK293T cells expressing HK24 or HK44 was stained by FITC-conjugated epitope-76 (60.7% and 40.8%, respectively) than HEK293T cells that did not express HK24 and HK44 (0.128%).

**Figure 2.**
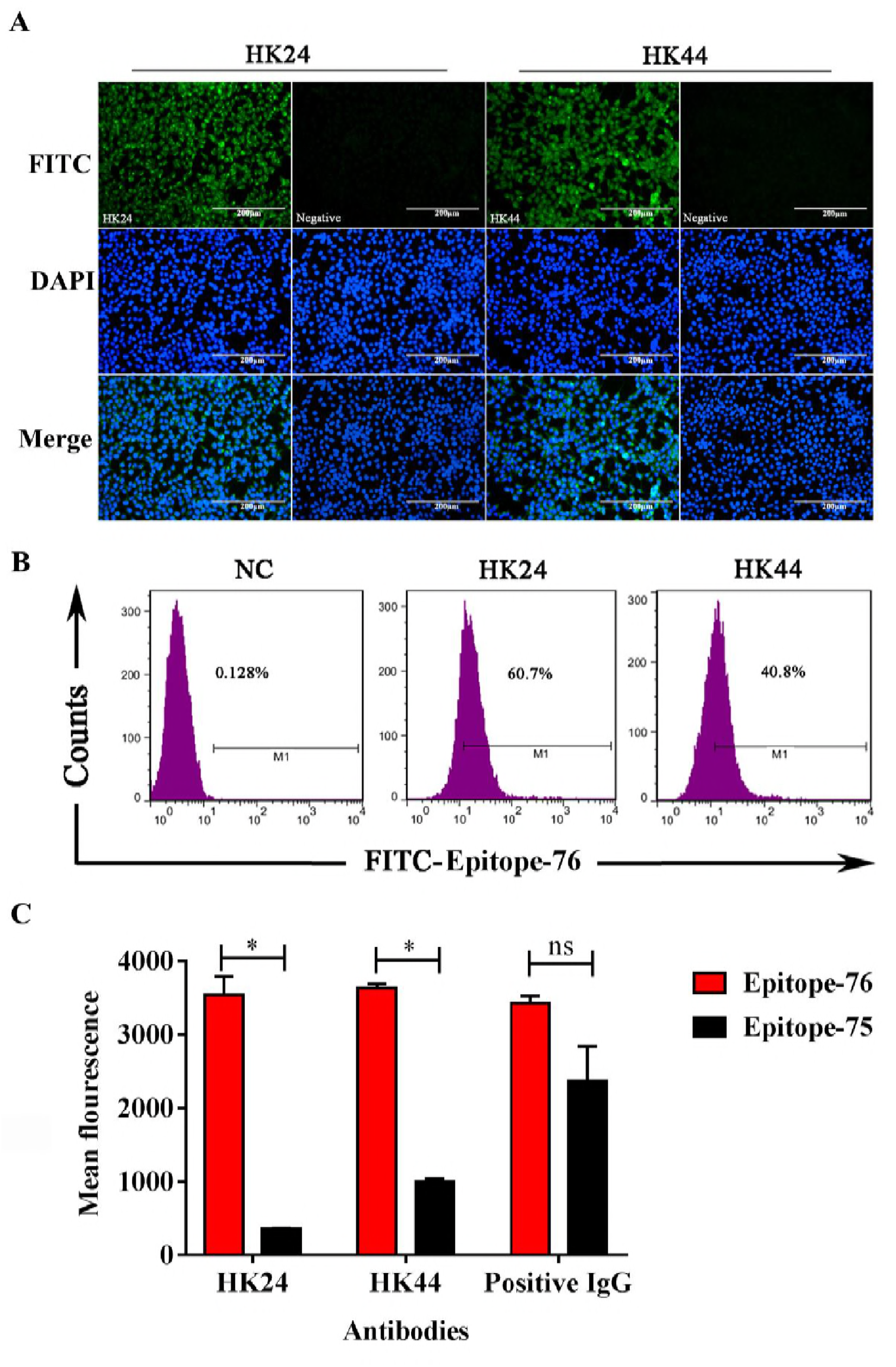
Specific reactions of the mAbs HK24 and HK44 with epitope-76. (A) Fluorescence microscopic images of cells. Green fluorescence was only observed on cells transfected with the HK24 or HK44 expression vectors. Cells transfected with empty vectors served as a negative control. Scale bar, 200 μm. (B) Analysis of the affinity of HK24 and HK44 for epitope-76. Affinity was analysed using FACS. NC is cells transfected with empty vectors. (C) Antibody specificity against epitope-76 was measured via a fluorescence assay. The mAbs HK24 and HK44 and positive IgGs isolated from pig #3748 were immobilized on an ELISA plate. Fluoresceinated epitopes were used to assess the binding activity of these antibodies. Epitope-75 is an unrelated epitope. The results from at least three biological replicates (mean ± SD) and analysed using t tests with GraphPad Prism software. p < 0.05, **p < 0.005, and***p < 0.0001.

We also performed a direct fluorescence experiment to ascertain whether the mAbs HK24 or HK44 possess binding specificity to epitope-76. As shown in **Fig. 2C**, the mAbs HK24 and HK44, as well as positive IgGs purified from pig #3748, were recognized efficiently by epitope-76, while HK24 and HK44 did not bind to the unrelated epitope-75 which is a linear epitope on the E2 protein of CSFV. Meanwhile, the positive IgGs interacted efficiently with both epitope-76 and epitope-75.

In summary, these data demonstrated that the mAbs HK24 and HK44 can be specifically recognized by epitope-76 with a high affinity.

### The mAbs HK24 and HK44 exhibit high specificity and sensitivity to the CSFV E2 protein

The reactivity between the mAbs and the CSFV E2 protein was deciphered via Western blotting. The results showed that the 55-kDa E2 protein band from lysates of CSFV-infected cells was clearly detected by the mAbs HK24 and HK44 with high specificity, unlike the lysates of uninfected cells (**Fig. 3A and 3B**).

**Figure 3.**
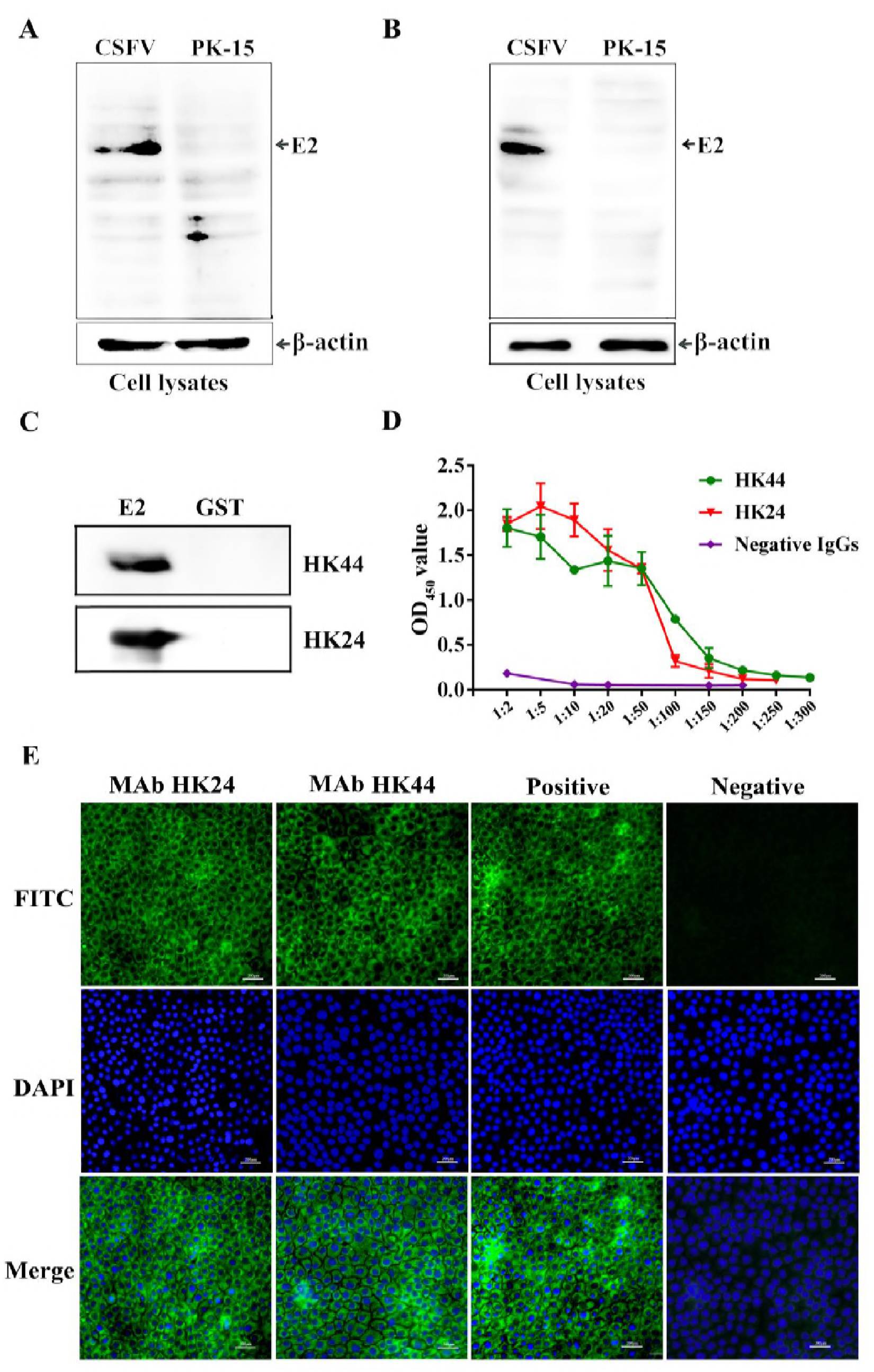
The mAbs HK24 and HK44 bound to the E2 protein. (A and B) The mAbs HK24 and HK44 bound to the CSFV E2 protein. Protein samples were obtained from PK-15 cells infected with CSFV. Uninfected PK-15 cells served as negative controls. All samples were resolved via 12% reducing SDS-PAGE and were identified using the mAbs HK24 (A) and HK44 (B). β-Actin was used as an internal control. (C) Recombinant CSFV E2 protein (690-866 aa) expressed in a bacterial system was recognized by the mAbs HK24 and HK44 and determined via Western blotting. (D) Detection of the binding of the mAbs HK24 and HK44 to the recombinant E2 protein using ELISA. Recombinant E2 protein (2 μg/ml) was used to coat the ELISA plates. The mAbs HK24 or HK44 (200 μg/ml), which were serially diluted in PBS, were added to each well. Negative IgGs were purified from CSFV-negative serum and used as a negative control. Results were obtained from at least three biological replicates (mean ± SD). (E) Detection of CSFV by the mAbs HK24 and HK44. PK-15 cells were infected with CSFV and identified via IFA using HK24, HK44, anti-CSFV swine serum and negative swine serum. Green fluorescence was observed on cells treated with HK24, HK44 and anti-CSFV swine serum. However, cells treated with negative serum showed no fluorescence. Scale bar, 200 μm.

To further verify the reactivity of the mAbs HK24 and HK44 with the CSFV E2 protein, recombinant E2 protein was subsequently produced and used to detect antibody affinity via Western blotting. As expected, the mAbs HK24 and HK44 reacted with recombinant E2 with high specificity (**Fig. 3C**). Indirect ELISA also showed that the mAbs HK24 and HK44 exhibited excellent affinity for the recombinant E2 protein, and binding occurred in a dose-dependent manner, whereas negative IgGs (isolated from CSFV-negative swine serum by protein A) did not show significant reactivity (**Fig. 3D**). Together, these results demonstrated that the mAbs HK24 and HK44 possess specific reactivity with either CSFV E2 protein in cells or purified recombinant E2 protein.

Additionally, to investigate whether the mAbs HK24 and HK44 could be applied for CSFV detection, these antibodies were used to detect CSFV in PK-15 cells via an immunofluorescence assay. As shown in **Fig. 3E**, the mAbs HK24 and HK44 showed strong fluorescence at 30 nM, as observed for CSFV-positive serum, unlike CSFV-negative serum. This result suggested that both mAbs isolated in the present study could be used to detect CSFV. Hence, our results suggested that the mAbs HK24 and HK44 exhibited high sensitivity towards E2 of CSFV and can be used for the serological detection of CSFV infection or for research purposes.

### Evaluation of the binding affinity of mAbs HK44 and HK24 to E2 protein by SPR

SPR is a powerful technique that provides affinity and kinetic information for protein-protein interactions (24, 25). Thus, SPR was used to assess the ability of the mAbs HK44 and HK24 to bind the recombinant protein E2. For this purpose, the mAbs HK44 and HK24 were immobilized on CM5 sensor chips with a general 1:1 interaction; the results are shown in **Fig. 4**. As shown in **Fig. 4A and 4B**, SPR dose-dependent binding assays verified that the mAbs HK44 and HK24 bound the E2 protein with high affinity at E2 protein concentrations ranging from 125 nM to 2000 nM. The SPR data indicated that the association rate constant (ka) of the mAbs HK44 and HK24 with the recombinant E2 protein was 1.70 × 10^4^ M^−1^ s^−1^ and 5.01 × 10^4^ M^−1^ s^−1^, indicating a fast association constant between mAbs HK44 or HK24 and E2 protein and the equilibrium dissociation constant (KD) of 1.85 × 10^−7^ M and 3.67 ×10^−8^ M respectively. These results suggest that mAbs HK44 and HK24 exhibit a high binding affinity to the E2 protein. The responses of E2 protein at each concentration to mAb HK44 or HK24 are depicted as a sigmoidal dose response curve in Fig. 4 C and 4D. These results clearly illustrate the higher specific affinity of the mAbs HK44 and HK24 for the E2 protein.

**Figure 4.**
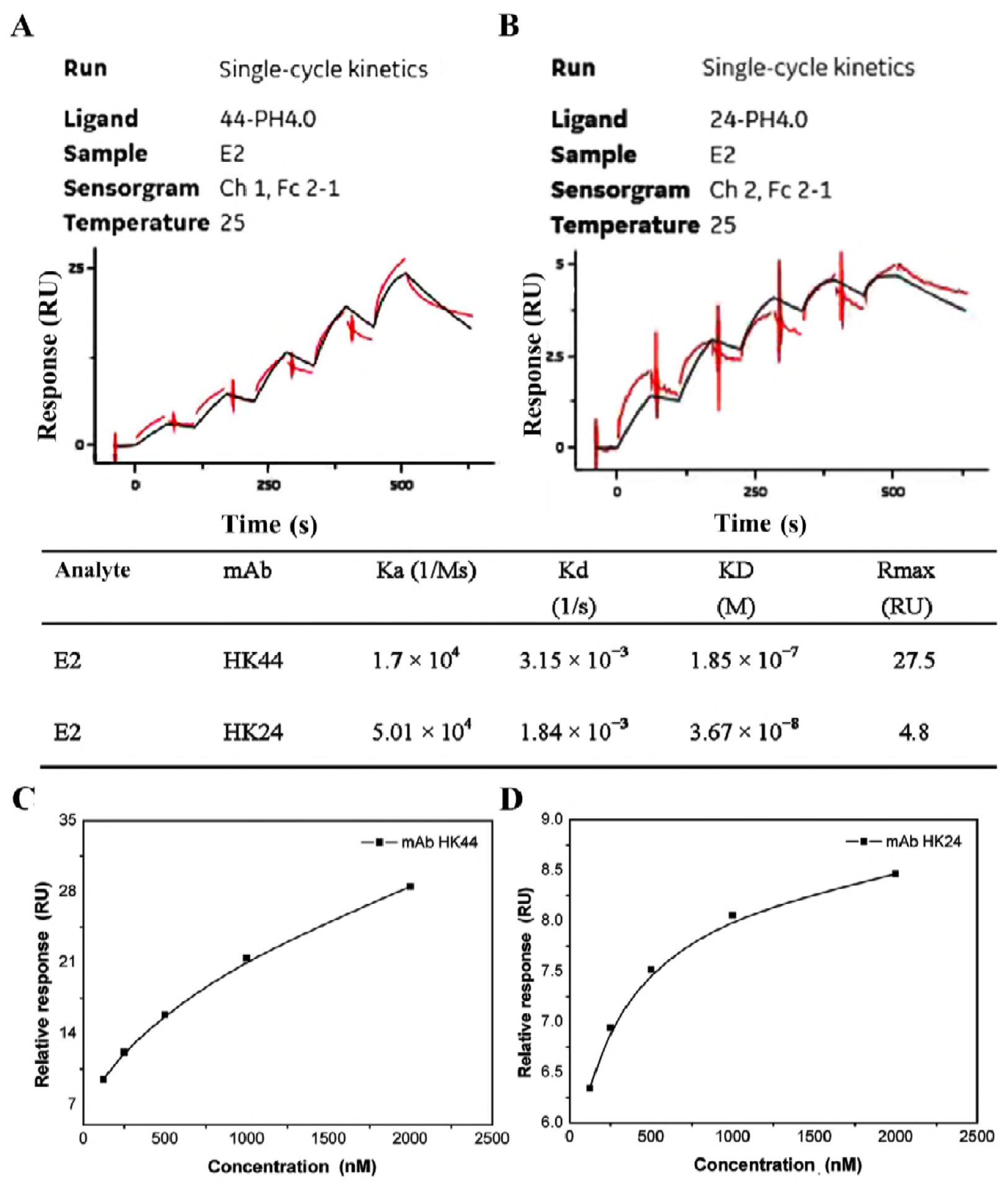
SPR binding assays. MAbs HK44 and HK24 bound to recombinant E2 protein. (A and B) Binding of E2 protein at concentrations of 2000, 1000, 500, 250 and 125 nM to mAbs (A) MAb HK44, (B) MAb HK24. The SPR data for each binding step were calculated and are shown in the table at the bottom. (C and D) Dose-response curve of the E2 binding signal. (C) The mAb HK44 binding response against each concentration of E2 protein. (D) The mAb HK24 binding response against each concentration of E2 protein.

### Identification of the binding of the mAbs HK24 and HK44 to a conserved epitope on the CSFV E2 protein based on a 3D structural model

To obtain detailed structural information on E2-mAbs interactions, protein-protein docking was performed. Before docking, the best protein modelling result was selected based on the probability density function (PDF) total energy and discrete optimized protein energy (DOPE). The PDF total energy and DOPE of HK24, HK44 and the E2 protein were 2296.5857, −45243.882812; 2390.8101, −46726.902344; and 1949.7089, −28135.984375, respectively. Furthermore, there were fewer irrational amino acid residues in the Ramachandran Plot, suggesting that the models we built may exist. Thus, protein-protein docking was performed by setting the Fab as the receptor protein and the E2 as the ligand protein. The docking results of E2-HK24 and E2-HK44 are shown in (**Fig. 5A and 5B**). More precisely, we found that the epitope Cys141-Lys155 of the E2 motif in both cases was buried in the middle of VL-VH interface of the E2-HK24 or E2-HK44 protein complex, and this epitope contacted the Fab CDR H3, CDR L3 and CDR L1 loops in both structures in addition to CDR H2 in the HK44-E2 complex (**Fig. 5C and 5D**). In addition, the mAb HK44 also directly bind to Thr88 and Pro90 (a linear epitope on E2 protein spanning Leu84-Pro90 residues that has been reported) (26, 27) by means of three residues (Val111A, Ser111B and Tyr111C) on the tip of the loop which is a relatively long CDR H3 loop (21 amino acids) (**Fig. 5C**). As shown in **Fig. 5D**, similar to E2-HK44 protein complex, the long CDR H3 of mAb HK24 (17 amino acids) not only binds to the epitope-76, but also binds to the linear epitope on E2 protein spanning Leu84-Pro90, via some hydrogen bonds and hydrophobic interaction. Detailed interactions between antigen and antibodies are shown in the **Table 2 and 3**.

**Figure 5.**
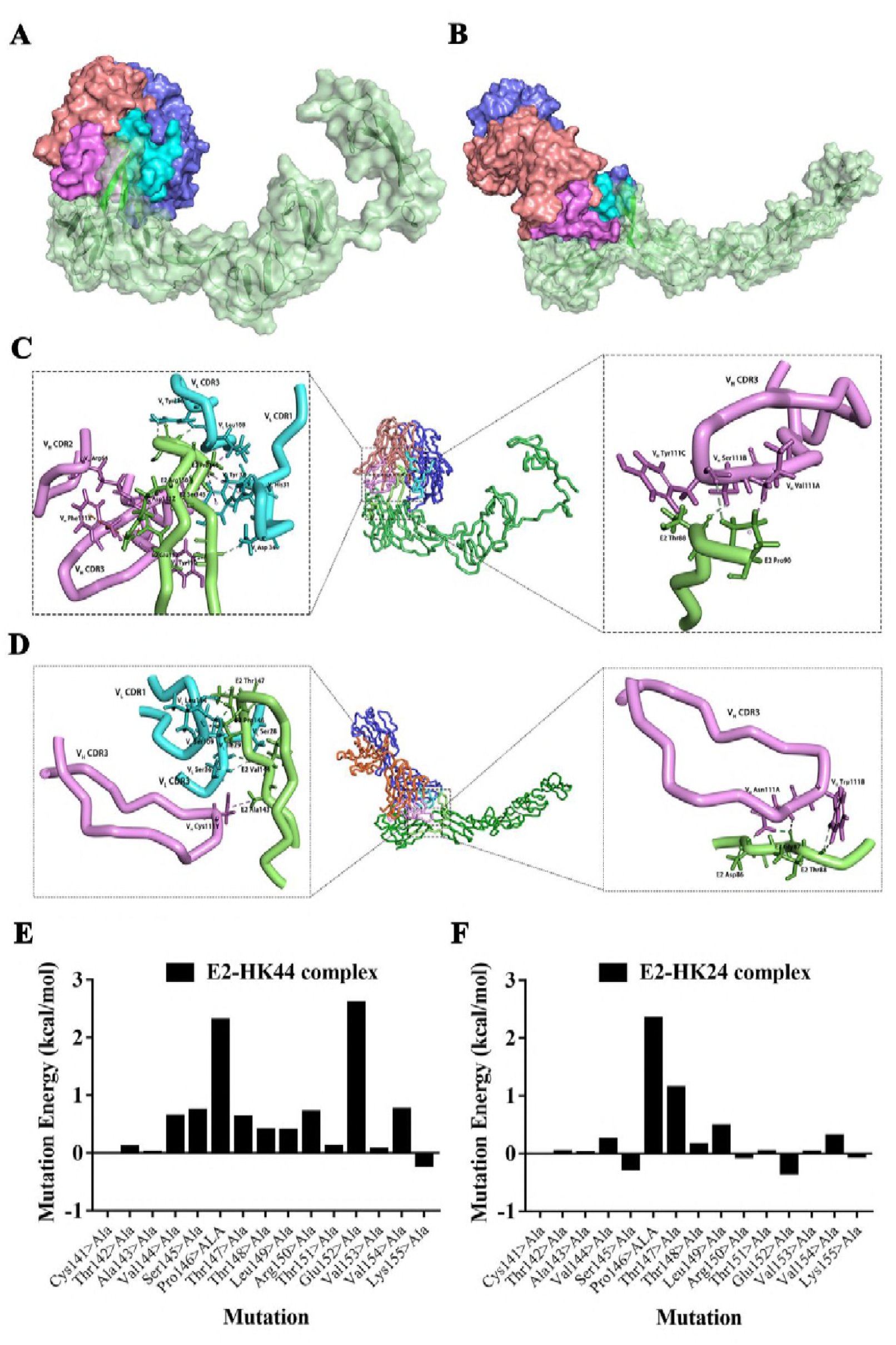
3D modelling of the combination of the Fab fragments of mAbs HK24 and HK44 with the CSFV E2 protein. (A and B) Cartoon representation of E2-HK44 and E2-HK24 protein complex. The best docking results are displayed by the surface. Light and heavy chains of the antibody are shown in slateblue and salmon, except for the CDR loops, which are shown in cyan and violet. The E2 protein is shown in pale green, except for the epitope, which is shown in green. (A) E2-HK44 protein complex. (B) E2-HK24 protein complex. (C and D) Bonds in the interface of the protein complex. Conventional hydrogen bond is colored in green, carbon hydrogen bond is colored in cyan, electrostatic (attractive charges and Pi-Cation) is colored in orange, Alkyl Hydrophobic is colored in pink, Pi-Sigma Hydrophobic is colored in purple. (C) E2-HK44 protein complex. (D) E2-HK24 protein complex. (E and F) The value of mutation energy during Virtual Alanine Mutation on the epitope-76 in the protein-protein complex. (E) E2-HK44 protein complex. (F) E2-HK24 protein complex.

To further confirm the interaction between mAb HK44 or HK24 and epitope-76, Virtual Alanine Mutation Scanning (VAMS) was employed to calculate mutation energy of individual amino acid residues changes using Discovery Studio 2017. In the case of E2-HK44 complex, when the residues of epitope-76 of E2 protein were individually mutated to alanine, seven residues on epitope-76, namely, Val144, Ser145, Pro146, Thr147, Arg150, Glu152, Val154 generated the mutation energy higher than 0.5 kcal/mol, indicating an unstable protein complex (**Fig. 5E**). The mutations of epitope-76 in the E2-HK24 complex procedure also destabilize the protein complex (**Fig. 5F**). When we mutate all of the epitopes to alanine, the mutation energy is 3.63 kcal/mol for E2-HK24 protein complex and 7.28 kcal/mol for E2-HK44 protein complex, indicating that the protein complex is an unstable state. In summary, these results suggested that the mAb HK24 or HK44 binds to the E2 protein by the recognition of the epitope-76 (CTAVSPTTLRTEVV).

### MAbs HK24 and HK44 can significantly neutralize CSFV *in vitro*

We next evaluated the protective efficacy of the mAbs HK24 and K44 against CSFV infection in PK-15 cells. A neutralization test was first performed via indirect immunofluorescence assay (IFA). As shown in **Fig. 6A**, the mAbs HK24 and HK44 displayed high neutralizing activity against CSFV, as no fluorescence was observed at the concentration of 75 μg/ml, and only weak fluorescence signals were detected at the concentration of 1.1 μg/ml, which was same as that observed for CSFV-positive serum. In contrast, strong fluorescence signals were detected in cells treated with CSFV-negative serum, indicating no inhibition of CSFV infection. The range of 50% inhibitory concentrations (IC_50_) to the non-neutralizing concentration for mAbs HK24 and HK44 is presented in **Fig. 6B**. These results indicated that the mAbs HK24 and HK44 exhibited neutralizing activity against CSFV, and they can prevent CSFV infections in cells.

**Figure 6.**
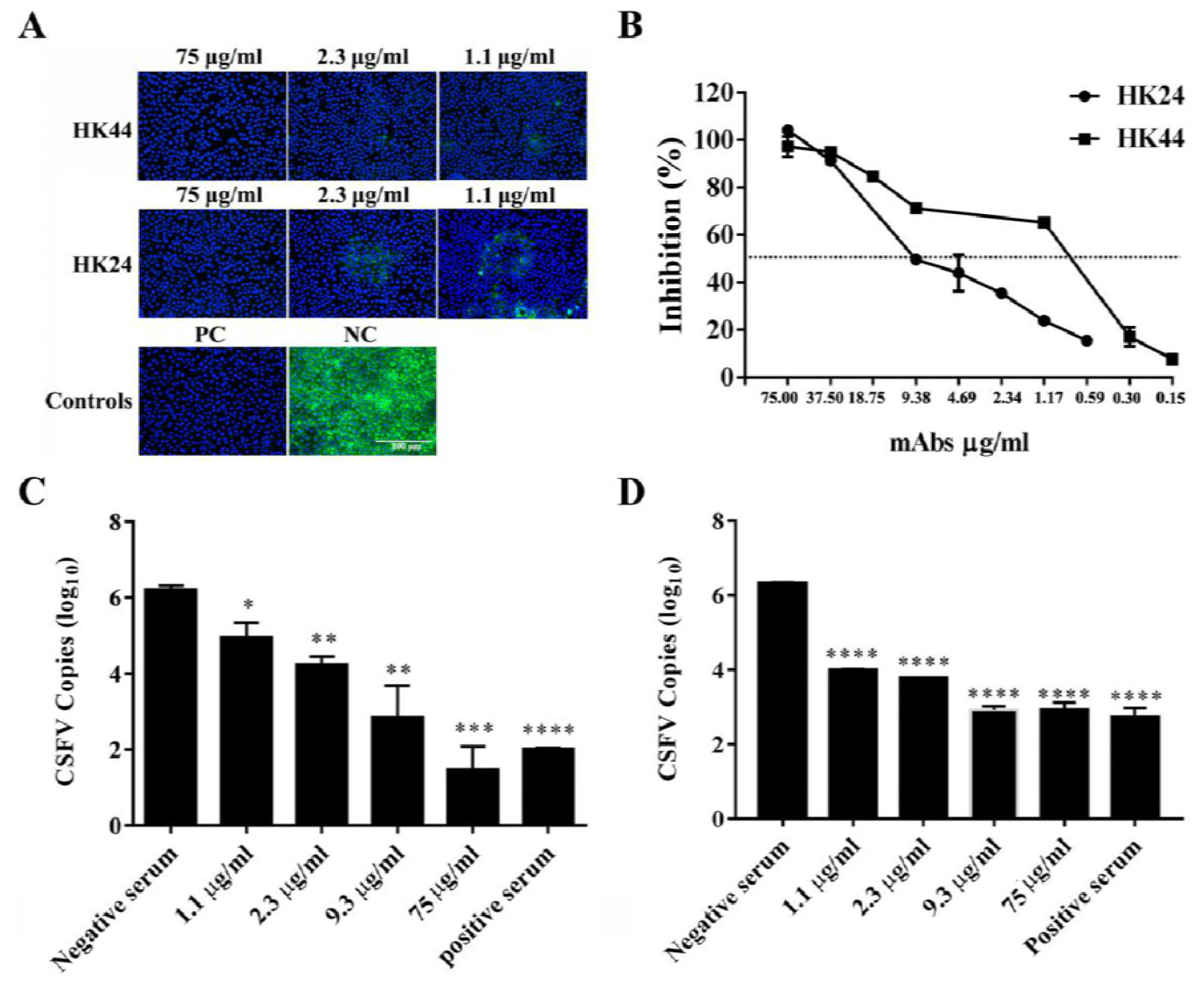
The mAbs HK24 and HK44 neutralize CSFV. (A) The mAbs HK24 and HK44 were tested for neutralization reactivity with CSFV via the IFA neutralization assay. CSFV-positive serum isolated from pig #3748 served as the positive control, and CSFV-negative serum isolated from pig #3741 was used as the negative control. (B) The percentage of CSFV inhibition. The dashed line represents 50% inhibition of the virus compared to the virus-only control. (C and D) Neutralization activity of the mAbs HK24 and HK44 was measured via quantitative qRT-PCR. (C) The number of CSFV copies in cells treated with the mAb HK24. (D) The number of CSFV copies in cells treated with the mAb HK44. Results were obtained from at least three biological replicates (mean ± SD) and analyzed using t tests with GraphPad Prism software. p < 0.05, p < 0.005, and p < 0.0001 (versus the positive serum).

To further directly detect whether the mAbs HK24 and HK44 block CSFV infection, the copy numbers of CSFV RNA in PK-15 cells were analysed via qRT-PCR with specific primers. The results suggested that treating PK-15 cells with the HK44-CSFV or HK24-CSFV complex at concentrations of 75 μg/ml to 1.1 μg/ml resulted in obviously lower numbers of virus copies in the cells than those observed with negative control treatment; at the same time, the mAb HK44 showed better neutralization activity than the mAb HK24 (**Fig. 6C and 6D**).

In summary, these data indicated that the mAbs HK24 and HK44 significantly neutralized CSFV and played an essential role in preventing CSFV from infecting target cells.

## Discussion

CSF is a disease that threatens the pork industry and has caused tremendous economic losses. As an alternative, development of effective vaccines and therapeutics agents may be a direct and effective approach to minimizing these losses. With the technological advances, mAbs now represent an important class of biotherapeutics, which can be used to treat autoimmune disorders diseases, cancer and viral or bacterial infections (28). Thus, it is necessary to establish a simple, highly efficient, low-cost method to produce anti-CSFV NAbs.

For generation of human mAbs, there have been previous reports of approaches involving the sorting of single B cells via FACS using fluorescently-labelled specific membrane immunoglobulins as antigen bait; However, this method may be limited by the number of antigenic baiting reagents (29–31). Here, we describe a method for efficient production of mAbs from immunized swine. We used a fluoresceinated linear neutralizing epitope CTAVSPTTLRTEVVK and FITC-labelled goat anti-pig IgG as bait for flow cytometry to screen B cells from a pig that had been pre-screened for high-level CSFV-specific antibody response in the blood. The cognate immunoglobulin heavy- and light-chain genes were then isolated from epitope-specific single B cells using single-cell RT-PCR technology. The key features of this method are the (a) sorting of B cells using a specific linear epitope by FACS, which may avoid the limitation of the availability of multiple antigenic baiting reagents; (b) direct isolation of natively matched full-length heavy- and light-chain pairs from a single B cell, thus, producing natural mAbs with a lower the risk of immunogenicity *in vivo* and greater suitability for *in vivo* applications; (c) rescue of mAbs that recognize the specific linear epitope. In contrast to traditional methods for generating mAbs, such as hybridoma technology, Epstein-Barr virus-transformation and phage-display libraries, this method helps to improve the recovery efficiency of mAbs as 78% of the mAbs produced showed neutralizing activity against CSFV. Taken together, this method is more convenient and efficient for obtaining epitope-specific NAbs against CSFV.

In this study, we amplified full-length heavy- and light-chain genes from one hundred single B cells and obtained nine mAbs, among which three were identical to HK24 and four to HK44. Thus, two mAbs HK24 and HK44 were selected to detect the strong binding reactivity against CSFV via Western blotting. To further define their binding paratopes, we tested the reactivity of the mAbs HK24 and HK44 with recombinant E2 via indirect ELISA and SPR. The results showed that the mAbs HK24 and HK44 exhibited comparable reactivity to the E2 protein. The binding sites within the linear epitope CTAVSPTTLRTEVVK on the E2 protein surface were predicted through a 3D structural model. Notably, these results are consistent with reports that the linear epitope CTAVSPTTLRTEVVK on the E2 protein is specifically recognized by CSFV antibodies (23, 32). Because the linear epitope is conserved in diverse CSFV strains and is divergent in the BVDV and BDV strains, we deduced that the mAbs HK44 and HK24 might be applied for the development of diagnostic reagents for CSFV.

We also confirmed that the mAbs HK24 and HK44 exhibited a powerful neutralizing response against the CSFV Shimen strain *in vitro*; in particular, the mAb HK44 showed an IC_50_ ≤ 1.1 μg/ml (**Fig. 6B**). This result indicated that these mAbs are capable of blocking virus infection and provided implications for therapeutic designs. Moreover, neutralization assays showed that in this work, the mAb HK44 exhibited more excellent neutralizing activity than the mAb HK24. We deduced that this may be due to a longer CDR H3 (21 amino acids) because the amino acid lengths of the predominant CDR H3 are conserved in most vertebrates, averaging 12-16 amino acids (33). Indeed, some longer CDR H3 loops usually play an important role in the adaptive immune response such as virus neutralization as described previously (34–36). Elongated CDR H3 can better overcome the structural barriers presented by antigens to confer protective functions.

In conclusion, this is the first report to describe the isolation of whole-porcine NAbs against the E2 protein of CSFV from single B cells of a vaccinated pig using a specific linear epitope. In this study, we characterized the functions of the mAbs HK24 and HK44 through a panel of assays. The results demonstrated that HK24 and HK44 displayed high sensitivity to CSFV, indicating that these mAbs have great potential for the detection and treatment of viral infections. In addition, we established a simple and rapid method for the isolation of specific B cells. The mAbs that we generated were derived directly from porcine B cells and are therefore safer and more efficient than mAbs produced in mice, rabbits or other species.

## Materials and methods

### Cell lines and virus

PK-15 cells (porcine kidney cell line, ATCC, CCL-33) and HEK293T cells (human embryonic kidney cell line, ATCC, Manassas, VA) were cultured in Dulbecco’s minimal essential medium (DMEM, Gibco, America) supplemented with 10% heat-inactivated foetal bovine serum (FBS, Gibco, America), at 37°C in 5% CO_2_. CSFV (strain Shimen) was gifts from Dr. Changchun Tu (Academy of Military Medical Sciences, Changchun, China).

### Single B cell sorting

Blood samples were isolated from a swine with a high-level CSFV-specific antibody response that received primary immunization with an attenuated vaccine strain of CSFV and subsequent boosting two times at one-week intervals with the same vaccine. Peripheral blood mononuclear cells (PBMCs) were isolated and suspended in phosphate-buffered saline (PBS). cells were stained for 30 min at 4°C using FITC-labelled goat anti-pig IgG (Sigma-Aldrich, USA) and 5-TAMRA-conjugated epitope-76, which is a conserved epitope (CTAVSPTTLRTEVVK) on the E2 protein of CSFV (synthesized by China Peptides Co., Ltd., China) (37). All buffers used for staining contained 2% FBS (vol/vol) to block non-specific binding. Single cells were sorted via flow cytometry into 0.2-ml thin-walled PCR tubes containing 20 μl of cell lysis buffer supplemented with 1 μl of RNase inhibitor, 19 μl of 0.2% (vol/vol) Triton X-100 (Sigma-Aldrich, USA), 1 μl of oligo-dT primer and 1 μl of dNTP mix. The cells were stored at −80°C until use (38, 39).

### Isolation and expression of Ig gene

PCR was performed to amplify the full-length immunoglobulin heavy- and light-chain genes (IgH and *IgL*) from single B cells as described previously (40, 41). Briefly, to obtain Ig genes, we performed a reverse transcription reaction using SuperScript II reverse transcriptase (38, 42). Next, the *IgH, Igλ* and *Igκ* genes were amplified separately via nested PCR reactions using a mixture of primers that were specific for heavy chain, kappa light chain and lambda light chain; all oligonucleotide primers are listed in **Table 1**. The PCR products of the full-length heavy- and light-chain genes were confirmed via sequencing and were separately cloned into the pEGFP-C1 expression vector with the EF1α promoter and WPRE element (**Fig. S1**). To produce mAbs, plasmids encoding immunoglobulin heavy- and light-chains were transiently co-transfected into HEK293T cells using PowerTrans293 Transfection Reagent (Throne Science, Shanghai, China) following the manufacturer’s instructions. After culturing for four days at 37°C under 5% CO2 in 6-well plates, supernatants were harvested via centrifugation, and antibodies were purified using Protein A (GenScript, USA Inc.) and quantified using a NanoDrop 2000.

**TABLE 1.**
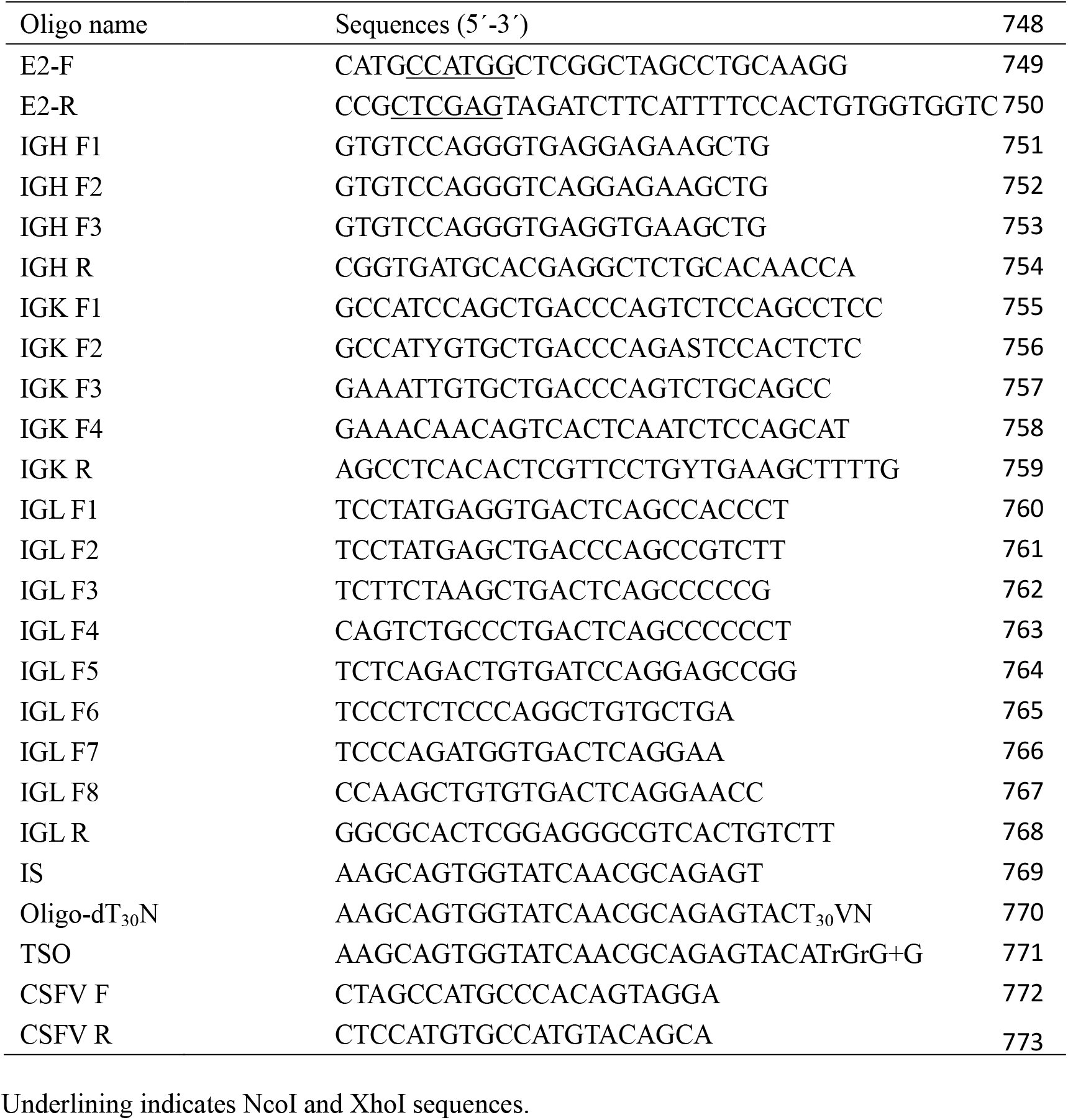
Oligoes used in this study.

**TABLE 2.**
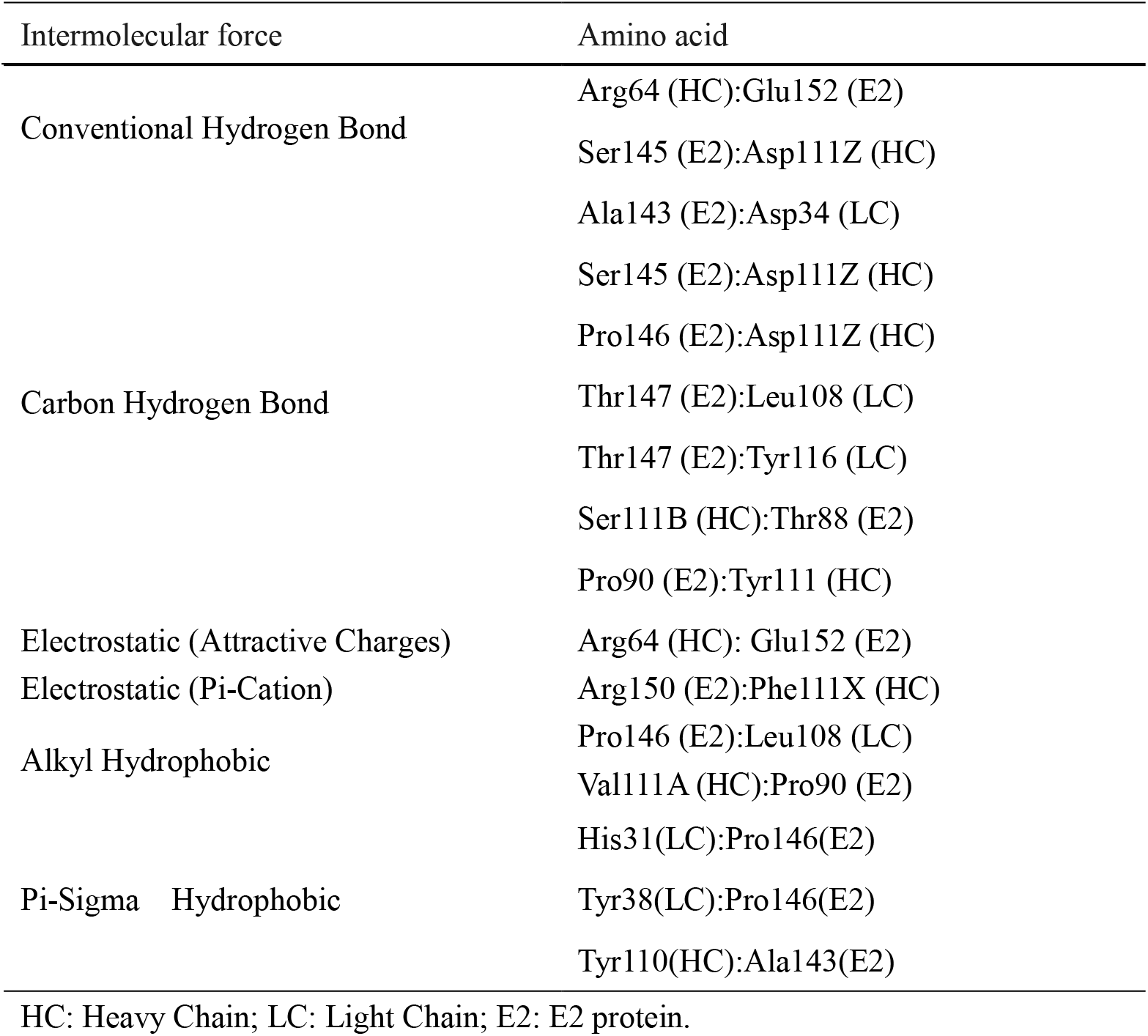
Intermolecular force affecting the interaction of mAb HK44 and E2 proteins.

**TABLE 3.**
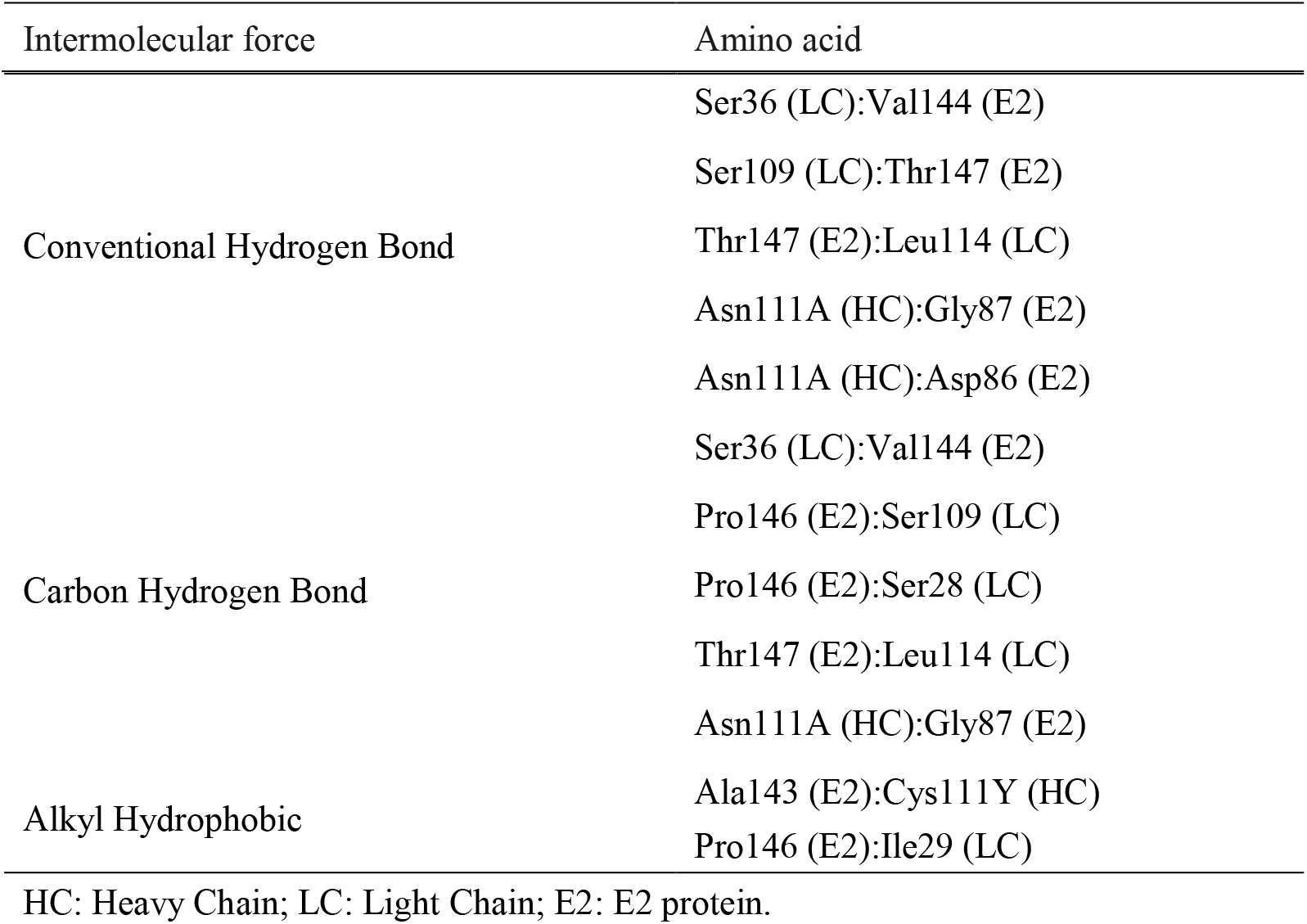
Intermolecular force affecting the interaction of mAb HK24 and E2 proteins.

### SDS-polyacrylamide gel electrophoresis (SDS-PAGE) and Western blotting analysis

SDS-PAGE was performed as follows. Briefly, antibody samples were separated in 12% (reducing conditions) resolving gels and 4% stacking gels, and protein bands were stained using Coomassie brilliant blue R-250 (43). For Western blotting analysis, we collected CSFV-infected cells and uninfected cells separately in cold PBS and lysed them using a lysis buffer (Beyotime, China) that contained 1 mM phenylmethylsulfonyl fluoride (PMSF) for 5 min on ice. After centrifugation at 10,000 × g at 4°C for 10 min, the supernatants were harvested and transferred into new 1.5-ml centrifuge tubes (44). Protein concentrations were determined with an enhanced-bicinchoninic acid (BCA) protein assay kit (Beyotime, China).

Western blotting was performed according to a previously described procedure (45). Briefly, equal amounts of proteins were loaded on a 12% SDS-PAGE gel. Next, the separated proteins were transferred onto nitrocellulose membranes, which were blocked with 5% (wt/vol) non-fat milk in Tris-buffered saline that contained 0.2-0.4% Tween-20 (TBS-T) for 2 h at room temperature. The membranes were probed with 4 μg/ml mAb or porcine IgGs (hyperimmune serum). Next, the bound antibody was stained using 2 μg/ml peroxidase (HRP)-conjugated rabbit anti-pig antibodies (Beijing Biosynthesis Biotechnology Co. Ltd, China). Immunoreactive bands were visualized using the ECL detection reagent (Beyotime, China) and the Azure c600 Western blot imaging system.

### Immunofluorescence to detect the binding specificity of mAbs HK44 and HK24 to epitope-76

The binding of epitope-76 to mAbs HK44 and HK24 were determined via immunofluorescence. In this procedure, enzyme-linked immunosorbent assay (ELISA) plates were coated with 100 μl (2 μg/ml) of a positive polyclonal antibody, mAbs HK24 and HK44 diluted in carbonate bicarbonate buffer (pH 9.6) and were incubated at 4°C overnight (46). The next day, the plates were washed thrice with PBS containing 0.05% Tween-20 (PBS-T), and 100 μl of blocking buffer (5% non-fat dry milk in PBS) was added per well. The plates were incubated at 37°C for 1 h. After blocking, 100 μl of FITC-conjugated epitope-76 or unrelated FITC-conjugated epitope-75 was added into each well, and the plates were incubated at 37°C for 2 h. Finally, fluorescence intensity was measured using a Tecan Microplate reader.

### Expression of the E2 protein

The partial sequence of E2 gene (GenBank: AY775178.2) was amplified from CSFV-infected cell mRNA via PCR and cloned into an expression vector (47). The vector was transformed into *E. coli* Rosetta (DE3) cells, and the transformants were selected and cultured in LB. The overnight cultures were subcultured at 1:100 and grown to an OD_600_ of 0.6 at 37°C. Next, 0.1 mM isopropyl-b-D-thiogalactopyranoside (IPTG) was added to induce the expression of the protein at 25°C for 12 h. The induced culture was harvested via centrifugation at 6,500 ×g for 15 min at 4°C and sonicated on ice. Subsequently, the inclusion bodies were separated from the crude cell lysate via centrifugation at 10,000 × g for 10 min and dissolved in 6 M urea at room temperature for 6 h. Finally, soluble E2 protein was purified through Ni-NTA resin as described previously (48, 49).

### Detecting the affinity of mAbs HK24 and HK44 for the CSFV E2 protein via indirect ELISA

An indirect ELISA was used to determine the affinity of the mAbs for the CSFV E2 protein. Purified E2 protein (2 μg/ml) was used to coat ELISA plates at 4°C overnight. The next day, the plates were washed thrice with PBS-T, and 100 μl of blocking buffer (5% nonfat dry milk in PBS) was added per well. The plates were incubated at 37°C for 1 h. After blocking, 100 μl of antibody samples were added per well, and the plates were incubated at 37°C for 2 h. Thereafter, an HRP-conjugated rabbit anti-pig antibody (Beijing Biosynthesis Biotechnology Co. Ltd, China) was added into each well at 37°C for 1 h. Finally, the wells were washed and incubated with 100 μl of TMB/H_2_O_2_ substrate per well at room temperature for 10 min. The reaction was stopped using 50 μl of 2 M H_2_SO_4_ per well, and the absorbance was read at 450 nm.

### Modelling of the E2 protein and the antigen-binding (Fab) fragment of the mAbs HK24 and HK44

The interactions of mAbs HK24 and HK44 with the CSFV E2 protein were simulated using Discovery Studio 2017. To obtain a three-dimensional model of the mAb HK24, we searched for a similar sequence in the Protein Data Bank (PDB) database. We found that the sequence homology between an anti-G-quadruplex-containing RNA antibody crystal (PDB 4KZD) and Fab of HK24 consisted of 84.1% similarity and 72.7% identity. Therefore, we used PDB 4KZD as a template for modelling the mAb HK24 Fab. Considering the quality of mAb HK44 model, we selected a chimeric template for modelling. For this purpose, humanized antibody 4B12 Fab (PDB 4LKX), an anti-TDRD3 FAB (PDB 3PNW) and an anti-CMV Fab Fragment (PDB 4LRI) were used as templates for modelling the light chain (LC), heavy chain (HC) and the interface of the mAb HK44 Fab; the sequence homologies (similarity and identity) between each template and the mAb HK44 Fab were 87.7% and 74.9% for V_L_, 80.4% and 70.9% for V_H_ and 82.8% and 69.8% for the entire Fab, respectively. E2 protein modelling was performed using the template of the crystal of the BVDV1 envelope glycoprotein E2 (PDB 2YQ2, 60% sequence identity with E2) (50).

### Docking and analysis of protein complexes

Before docking, models were typed using the CHARMm Polar H forcefield. We set E2 as the ligand protein and the Fab of HK24/HK44 as the receptor protein. ZDOCK, a rigid-body protein-protein docking algorithm based on the Fast Fourier Transform Correlation technique, was used with a 6° angular step size to generate 54,000 poses, of which the top 2000 were re-ranked by ZRANK (51). These poses were then processed with RDOCK, and only the top 10 clusters with the highest density of poses were further considered. Finally, we used the “Analyze Protein Interface” protocol for analysis. All docking calculations were performed using Discovery Studio 2017.

### Surface plasmon resonance (SPR) analysis

To analyse the affinity of the mAbs for the CSFV E2 protein, we performed SPR analysis on a Biacore 8K system (GE Healthcare) using CM5 sensor chips (GE Healthcare) and PBS-T as the running buffer during immobilization and binding analysis at a constant temperature of 25°C (52). Initially, the surface was activated with 1-ethyl-3-(3-dimethylaminopropyl)-carbodiimide (EDC) and N-hydroxysuccinimide (NHS), and the mAb HK44 or HK24 was diluted in 10 mM sodium acetate buffer (pH 4.0) and immobilized on the CM5 sensor chip using Amine Coupling Kit (GE Healthcare). Next, for binding analysis, five different concentrations of the recombinant E2 protein diluted in the running buffer (2000 nM, 1000 nM, 500 nM, 250 nM, and 125 nM) were allowed to flow over the chip surface. After sample injection was completed, the running buffer was allowed to flow over the surface to perform dissociation. At the end of dissociation, the sensor surface was regenerated with a glycine-HCl solution (53). Finally, all experimental data were analysed using Biacore 8K evaluation software (GE Healthcare).

### Neutralization assays

The neutralization activity of the mAbs HK24 and HK44 against CSFV was tested via an immunofluorescence assay as described previously (54). Briefly, PK-15 cells were grown to 30-40% confluence in complete DMEM containing 10% FBS on 96-well plates at 37°C in 5% CO_2_. Antibodies (150 μg/ml) were diluted two-fold in DMEM in a series and mixed 1:1 with 100 TCID_50_ CSFV, and the mixtures were incubated at 37°C for 1 h. After incubation, 100 μl of the antibody-virus mixture was added into the wells of the 96-well plates to infect the PK-15 cells for 2 h. Uninfected cells and virus-infected cells acted as positive and negative controls, respectively. After 2 h of incubation, the supernatants were aspirated, and the cells were washed three times with PBS. Next, 200 μl of fresh medium was added, and the cells were incubated at 37°C for 72 h. Subsequently, the cells were fixed with 80% cold acetone at -20°C overnight. The next day, the cells were washed and incubated with porcine CSFV-positive serum (1:100 dilution in PBS containing 10% FBS) at 37°C for 2 h. After being washed three times with PBS, FITC-labelled goat anti-pig IgG (Sigma-Aldrich, America) was added into each well, and the plates were incubated at 37°C for 0.5 h (55). Images were obtained using fluorescence microscopy (Olympus BX51).

### Quantification of CSFV RNA

Quantitative RT-PCR (qRT-PCR) was performed to examine CSFV in PK-15 cells. Total RNA was extracted from the uninfected and virus-infected cells using TRIzol (Tiangen, Beijing, China), according to the manufacturer’s instructions. Reverse transcription reactions were performed using the FastKing RT Kit (Tiangen, Beijing, China) to synthetize cDNA, and qRT-PCR was performed on a Bio-RadiQ5 instrument (BioRad, USA) with SuperReal PreMix Plus (Tiangen, Beijing, China). A standard curve was simultaneously created to calculate the viral load in each sample.

### Statistical analysis

All data presented in the figures are expressed as the mean ± SD from at least three independent experiments. When the data from the neutralization assays were analysed, the antibody concentrations were transformed to log_10_, and the IC_50_ was calculated using GraphPad Prism software 7.0 (La Jolla, CA, America) with the equation for dose response (variable slope). P < 0.05 was considered statistically significant.

## Acknowledgements

We would like to thank Xue Chen for technical assistance. This study was financially supported by a grant from the Special Funds for Cultivation and Breeding of New Transgenic Organisms (No. 2016ZX08006003), the Program for JLU Science and Technology Innovative Research Team (2017TD-28), the Program for Changjiang Scholars and Innovative Research Team in University in China (No. IRT16R32), the Fundamental Research Funds for the central Universities, the Jilin Province Science and Technology Development Projects (No. 20150204077NY) and the National Natural Science Foundation of China (No. 31772747).

## Conflict of Interest

The authors declare that they have no competing interests.

## Ethical Approval

Not applicable.

---

**Figure S1.**
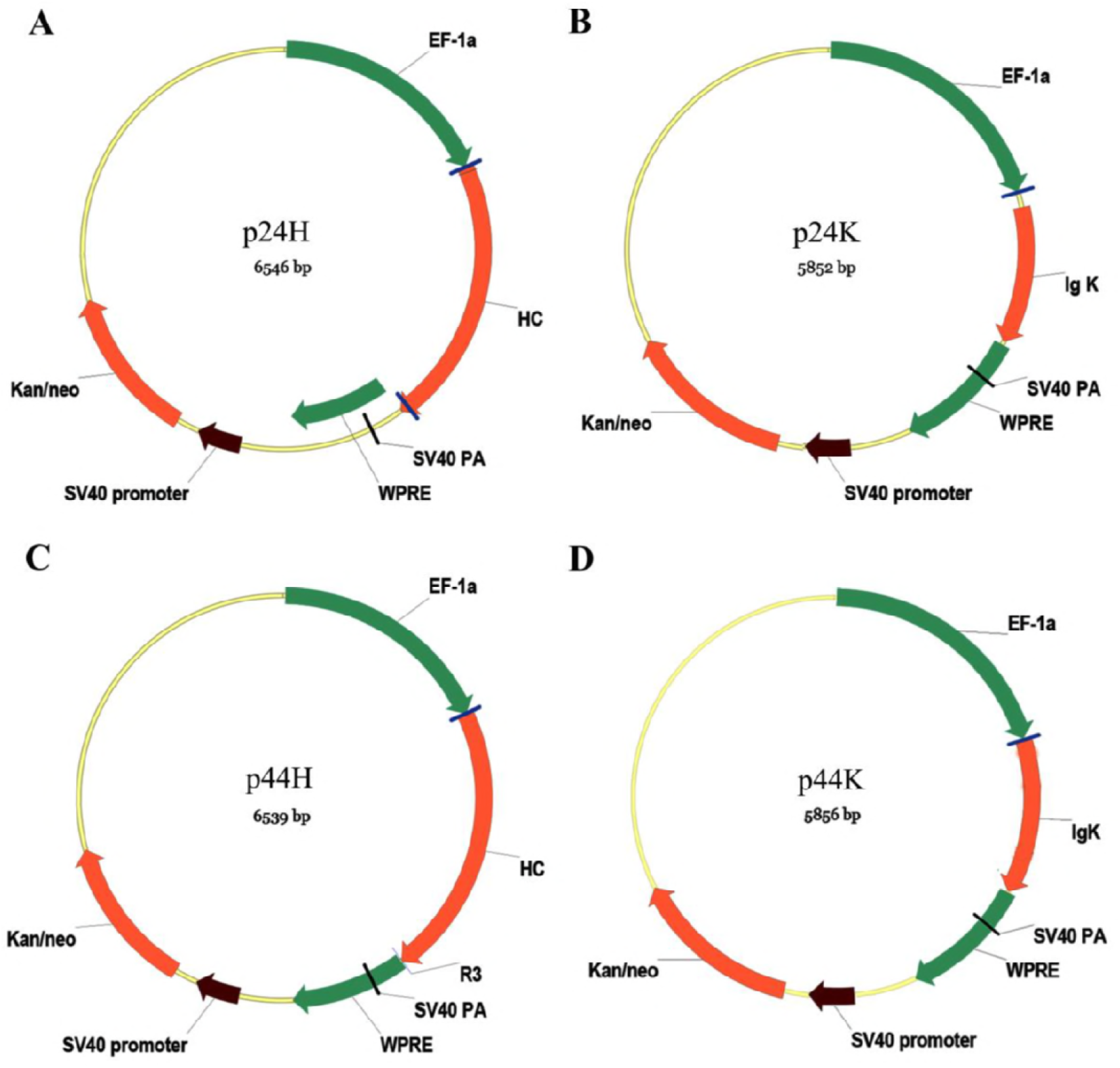
Physical maps of the expression vectors of the mAbs HK24 and HK44. Plasmid pEGFP-C1 was selected as the backbone, and the CMV promoter was substituted by the EF1α promoter. (A) P24H is the expression vector of the kappa light chains of HK24. (B) P24K is the expression vector of the heavy chains of HK24. (C) P44H is the expression vector of the kappa light chains of HK44. (D) P44K is the expression vector of the heavy chains of HK44.

